# Patterns and Causes of Signed Linkage Disequilibria in Flies and Plants

**DOI:** 10.1101/2020.11.25.399030

**Authors:** George Sandler, Stephen I. Wright, Aneil F. Agrawal

**Affiliations:** Department of Ecology and Evolutionary Biology, University of Toronto, 25 Willcocks Street, Toronto, ON M5S 3B2, Canada; Center for Analysis of Genome Evolution and Function, University of Toronto, 25 Willcocks Street, Toronto, ON M5S 3B2, Canada

## Abstract

Most empirical studies of linkage disequilibrium (LD) study its magnitude, ignoring its sign. Here, we examine patterns of signed LD in two population genomic datasets, one from *Capsella grandiflora* and one from *Drosophila melanogaster.* We consider how processes such as drift, admixture, Hill-Robertson interference, and epistasis may contribute to these patterns. We report that most types of mutations exhibit positive LD, particularly, if they are predicted to be less deleterious. We show with simulations that this pattern arises easily in a model of admixture or distance biased mating, and that genome-wide differences across site types are generally expected due to differences in the strength of purifying selection even in the absence of epistasis. We further explore how signed LD decays on a finer scale, showing that loss of function mutations exhibit particularly positive LD across short distances, a pattern consistent with intragenic antagonistic epistasis. Controlling for genomic distance, signed LD in *C. grandiflora* decays faster within genes, compared to between genes, likely a by-product of frequent recombination in gene promoters known to occur in plant genomes. Finally, we use information from published biological networks to explore whether there is evidence for negative synergistic epistasis between interacting radical missense mutations. In *D. melanogaster* networks, we find a modest but significant enrichment of negative LD, consistent with the possibility of intra-network negative synergistic epistasis.

## Introduction

Linkage disequilibrium (LD), the association of different alleles across the genome, is a general feature of population genomic datasets, often revealing clues of ongoing evolutionary or demographic processes (McEvoy et al. 2011). For example, in finite populations, drift can be a ready source of LD, generating both positive and negative associations between alleles (Hill and Robertson 1968). While unsigned LD has been extensively studied in population genetics (through statistics such as r^2^), signed LD has received relatively less attention, despite the fact that the sign of allelic associations can also provide useful information. Here we refer to positive associations as those between two common alleles (or equivalently between two rare alleles), and negative associations as those between common and rare alleles. Demographic processes such as admixture and population structure can create LD, where unlike drift in a single panmictic population, an overabundance of positive associations is expected between pairs of migrant alleles (Chakraborty and Weiss 1988; Stephens et al. 1994; Pfaff et al. 2001). Selective processes can also be a source of LD; for example, ongoing strong selective sweeps can be characterised by an elevation of unsigned LD around the sweeping haplotype (McVean 2007). Non-independence of mutational events, e.g. multinucleotide mutations, arise at non-negligible frequencies in several species (Schrider et al. 2011), and could also be an important source of positive LD among de-novo mutations (Ragsdale 2021). Finally, unsigned LD can also be used to analyze patterns of recombination across the genome, as recombination is expected to break down any existing LD (Auton and McVean 2007).

LD can also build up due to selection against deleterious mutations in two different ways. First, Hill Robertson interference (resulting from the interaction of selection and drift) can cause negative associations to build up among deleterious mutations, if recombination between them is limited (Hill and Robertson 1966). In sexually reproducing organisms such as humans, this process has recently been suggested to build up negative LD among physically proximal, missense mutations (Garcia and Lohmueller 2020). Second, negative selection can cause LD among deleterious mutations to build up if epistasis is present (Kondrashov 1995; Sohail et al. 2017). Under the null model of multiplicative fitness, where each mutation contributes to a reduction in fitness independently of other mutations, LD is not expected to accumulate. Synergistic epistasis, where each additional deleterious mutation reduces fitness by a greater magnitude, creates negative LD among deleterious mutations and vice versa for antagonistic epistasis (Kimura and Maruyama 1966; Kondrashov 1982).

Synergistic epistasis among deleterious mutations is of particular interest because such epistasis has several evolutionary consequences. For example, negative synergistic epistasis allows for lower mutation loads under mutation-selection balance, and can influence the evolution of sex and recombination (Kimura and Maruyama 1966; Crow and Kimura 1970; Crow and Kimura 1979; Kondrashov 1982; see also Barton 1995). Despite considerable interest, empirical data on epistasis among deleterious mutations is limited with most data coming from microorganisms assayed in a lab setting. These studies have found that synergistic and antagonistic interactions are both common so that mean epistasis is close to zero (Elena and Lenski 1997; Agrawal and Whitlock 2010; Lalić and Elena 2012; Bank et al. 2015; Puchta et al. 2016). A recent study by Sohail et al. (2017) used a different approach to make inferences about epistasis. They examined patterns of signed LD among rare loss of function (LOF) mutations in humans and fruit flies demonstrating that across several datasets LOF mutations had significantly lower values of signed LD than their neutral reference (synonymous sites), a pattern consistent with the action of negative synergistic epistasis.

Here we examine patterns of LD across several classes of mutations in a published dataset of 182 individuals of *Capsella grandiflora* sampled from a population in Greece, and 191 *Drosophila melanogaster* flies sampled from an ancestral population in Zambia (Lack et al. 2015). We find that mean signed LD is positive for most types of mutations across the genome except for LOF mutations. The magnitude of positive LD scales with the predicted deleteriousness of the mutations we analyze, with more neutral mutations exhibiting the most positive LD. We use simulations to show that admixture or distance biased mating could produce this type of pattern and provide alternative explanations to epistasis for differences in LD among neutral versus deleterious mutations. We then explore finer scale patterns of LD and uncover strong short-scale positive LD among LOF mutations, a potential signal of within gene antagonistic epistasis. Further analyses show that within gene LD is generally stronger than between gene LD in *C. grandiflora* (correcting for distance between pairs of mutations), for both neutral and deleterious mutations. This pattern is broadly consistent with cross-over hotspots frequently occurring in promoter regions of plant genomes. Finally, we use gene network information from KEGG to explore signals of LD and epistasis among deleterious mutations segregating in functionally related genes. We report no significant LD in *C. grandiflora* but significantly more negative LD in *D. melanogaster* KEGG networks compared to a null distribution generated from permuted networks, a pattern that could indicate synergistic epistasis acting against gene flow.

## Results

We analyzed patterns of signed LD among several classes of mutations (synonymous, missense non-radical, missense radical, intronic, non-coding, UTR (untranslated region), and LOF), in a dataset of 182 outbred diploid *C. grandiflora* individuals, and a dataset of 191 *D. melanogaster* haploid embryos (population DPGP3). We only considered variants below a specified threshold for minor allele frequency because we hoped to maximize the probability that the rare variants at each site are deleterious, though the degree of that deleteriousness is expected to vary among mutational classes (e.g., for synonymous sites, rare variants are presumably negligibly more deleterious than the common variant on average). The sign of LD was polarized by frequency so positive/negative LD should indicate that deleterious variants are found more/less often together than expected.

We first measured mean LD by assessing the over- or under-dispersion of deleterious (or synonymous) variants among individual genomes (see Materials and Methods; (Sohail et al. 2017)). An under-dispersion of the deleterious variants implies negative LD (i.e., deleterious variants are found together less often than expected by chance). We calculated mean LD per pair of alleles using several different allelic count cut-offs (i.e., minor allele frequency thresholds). In both species, point estimates for mean LD were positive for all classes of mutations, and all allelic count cut-offs examined, except LOF mutations (Figure 1), where the point estimate for mean LD was negative using some allelic count cut-offs but not others. When repeating this analysis in *D. melanogaster,* excluding regions which were known to harbor inversions in the DPGP3 population, we found the same qualitative results albeit with slightly reduced positive LD for most mutations classes (Supplementary Figure 1).

**Figure 1.**
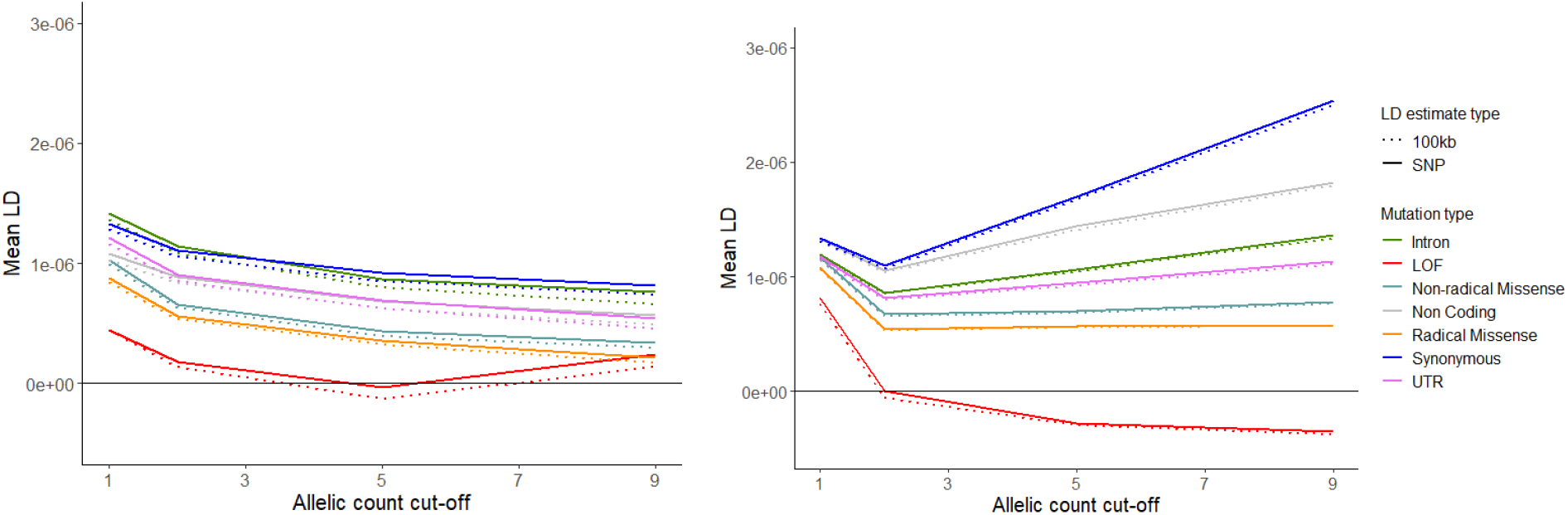
Mean pair-wise LD among several classes of mutations across different allele count cut-offs. Solid lines indicate mean LD among all SNPs, dashed lines indicate LD calculated among sites in different 100kb, non-overlapping genomic blocks. Left, results for *C. grandiflora*, right, results for *D. melanogaster.*

One pattern apparent in our data is that the least deleterious mutational classes exhibited the most positive mean LD in both flies and plants (i.e., the most positive LD belonged to classes such as intronic and synonymous). The site frequency spectra for these different mutational classes add support to the suspected rank ordering in the deleteriousness of different mutational classes (Supplementary Figure 2) such that the classes with the greatest excess of rare variants (presumed to be the most deleterious) had the more positive LD. In *D. melanogaster* the order of deleteriousness inferred from the site frequency spectra (starting with least deleterious) was as follows: synonymous, non-coding, intronic, UTR, missense non-radical, missense radical, LOF. Similarly, in *C. grandiflora* the order was synonymous, intronic, UTR, non-coding, missense non-radical, missense radical, LOF (Supplementary Figure 2).

The observation that LD was strongest for neutral/nearly neutral mutations suggests a non-selective force, such as admixture, is building LD (Sohail et al. 2017; but see also Good 2020). We used a series of simulations using SLiM (V3.2.1) (Haller and Messer 2019) to explore how different cases of non-equilibrium demography and population structure can affect LD among neutral and deleterious mutations (see Materials and Methods for more details). We first tested how a model of admixture might impact patterns of LD for rare neutral and deleterious mutations under strictly multiplicative selection. We simulated admixture between a focal population and two previously isolated satellite populations and polarized LD by variant rarity. We found that admixture easily caused positive LD to build up among neutral mutations, particularly so if admixture started recently between populations that had previously been isolated (Figure 2A). However, this was not the case for deleterious mutations in these populations, where LD remained much closer to 0 albeit slightly positive on average if admixture was present. This result was also apparent if we polarized LD by true ancestral state in our simulations and imposed a minor allele frequency cut-off, or if we polarized by frequency as in our real-world data, but did not implement a minor allele frequency cut-off (Supplementary Figure 3). The only case where we did not observe positive LD for neutral mutations was if we polarized LD by true ancestral state and implemented no allele frequency cut-off (Supplementary Figure 3).

**Figure 2.**
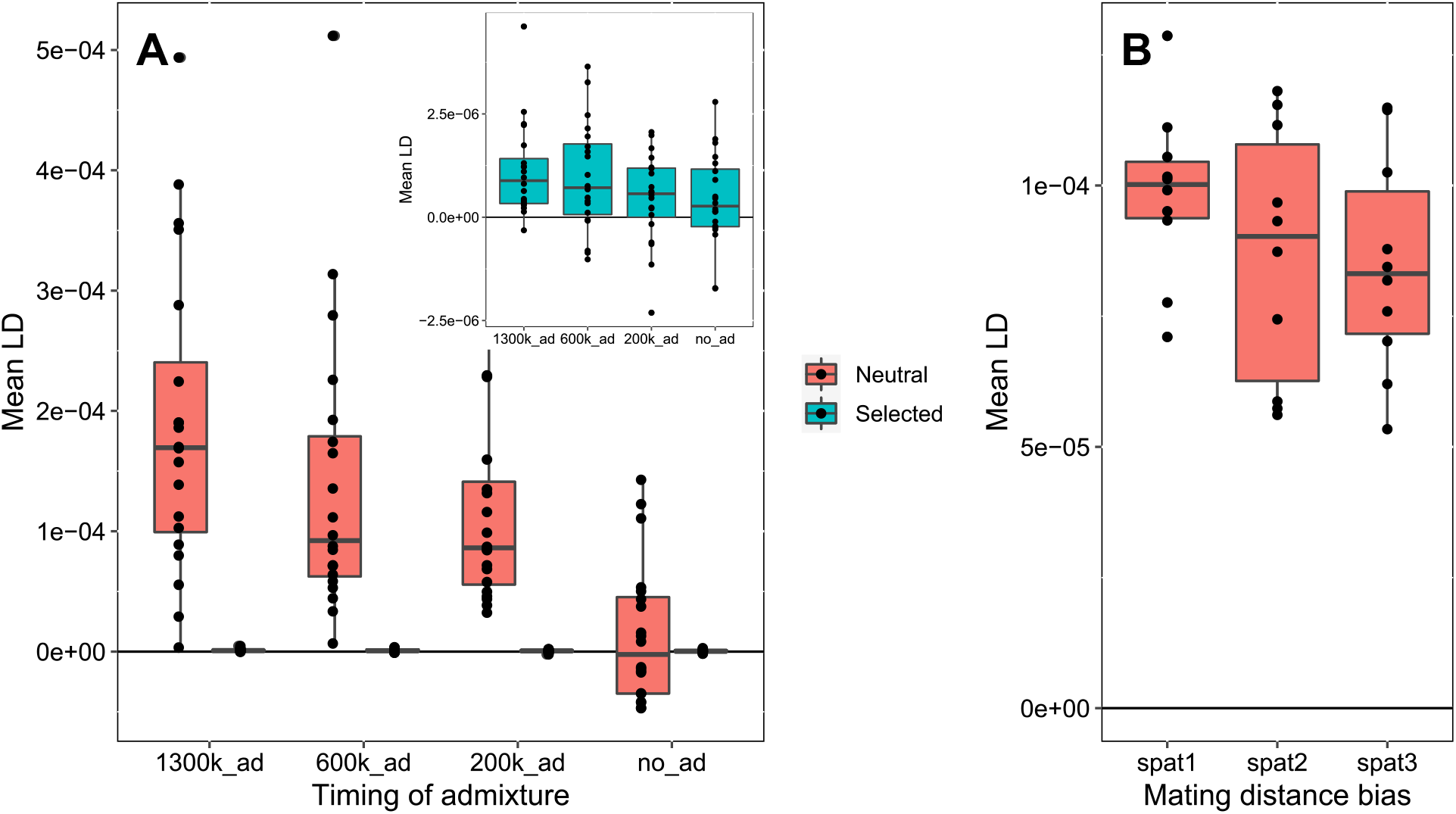
A) Mean signed LD among simulated neutral and deleterious mutations under different scenarios of admixture. The *x*-axis represents the generation in which admixture between isolated populations started. All simulations were run of a total of 1.5 million generations. Inset highlights results for selected (deleterious) mutations B) Mean LD among neutral mutations segregating in simulated populations existing on a 2D geographic landscape. The *x*-axis represents different scenarios of mating bias by distance with increasingly more random mating to the right of the *x*-axis.

We then explored isolation by distance due to continuous geography as a potentially common mechanism that could create positive LD in a similar way to admixture. Using SLiM to model populations on a continuous 2D landscape we again readily observed positive LD forming among neutral mutations under several scenarios of distance-biased mate choice, demonstrating that spatial considerations alone might be able to explain patterns of positive LD in our two datasets (Figure 2B).

Our simulation results qualitatively match the earlier simulation results reported by Sohail et al. who examined models specific to human demographic history (i.e., population structure and gene flow). They found positive LD does not build uniformly for deleterious and neutral mutations, rather, the more deleterious a class of mutations, the less positive LD built up among them. In summary, all of these simulations clearly show that spatial structure with gene flow or admixture creates a difference in LD for selected versus neutral sites.

While our patterns overall seem consistent with a relatively simple model of spatial structure with varying strengths of purifying selection across site types, some of our point estimates of LD for LOFs were negative, and negative LD is not expected under such models of gene flow. Rather negative LD could be indicative of synergistic epistasis or Hill-Robertson interference. To assess whether these processes might be creating negative LD in our datasets, we next tested whether our estimates of negative LD were significantly different from zero. We did this by permuting the assignment of LOFs among all individuals in each dataset. This method preserves the allele frequency at each locus while randomizing the associations among loci. We focused exclusively on LOF mutations at an allele count cut off of no more than 5 because this cut-off resulted in the most negative point estimates of mean LD in both datasets and such rare mutations are more likely to be truly deleterious. All our subsequent analyses utilize this allelic cut-off value for both datasets. This test suggested that LD among LOF mutations was not significantly different from 0 in either *C. grandiflora* or *D. melanogaster* when calculating LD SNP-by-SNP (p = 0.996 and p = 0.386, 2-tailed) or among sites in different 100kb blocks (p = 0.680 and p = 0.346, 2-tailed). When we applied this permutation approach to synonymous mutations, we found that LD was significantly greater than 0 in both species, when calculating LD SNP-by-SNP (p < 0.002 both species, 2-tailed), or using 100kb blocks (p < 0.002 both species, 2-tailed), further verifying positive LD among more neutral mutations. Again, removing regions with segregating inversions did not qualitatively change the results in *D. melanogaster* for LOF mutations (p = 0.658, p = 0.648, LD calculated SNP-by-SNP and using 100kb blocks respectively), or synonymous mutations (p < 0.002, for both types of LD estimates).

In the preceding sections, we examined genome-wide average LD. However, most pairs of sites contributing to this average are far apart or are found on different chromosomes. For such sites, meiotic recombination and segregation will very rapidly destroy any allelic associations formed by processes like selection. Significant signed LD however, could still be present between mutations that are physically proximal. We therefore next used PLINK (Purcell et al. 2007) to assess the relationship between inter-mutation distance and LD for each class of mutations. Consistent with our first analysis, LD was positive for all mutation classes in most distance bins; those estimates that were negative were small in magnitude and were not significantly different from zero (Figure 2). Within a distance bin, positive LD was stronger for the most weakly selected mutation classes for most distance bins.

An interesting exception to this pattern in both species was in the smallest distance bin (0-100bp). The major outlier in this distance bin were LOF mutations which had surprisingly positive mean LD estimates in both species. The confidence intervals on these estimates were very large for LOF mutations in this distance bin due to the small number of observations for the mutation class. However, the high LD estimate for LOFs is present in both species, and, in *C. grandiflora,* the 95% confidence intervals suggested LOF mutations had more positive LD than all other mutation types aside from intronic and synonymous. This pattern is consistent with intragenic antagonistic epistasis, which seems probable for true LOF mutations occurring within the same gene. Ideally, we would evaluate this hypothesis by comparing LD between physically close LOFs that occur in the same versus different genes. However, we had too few intergenic LOFs at short distances to do so.

Within gene antagonistic epistasis could also create positive LD among other types of deleterious mutations such as missense mutations, which are much more abundant. We compared signed LD decay within and between genes for both synonymous and non-radical missense mutations (the two coding classes with ample data) to test for this. In the case of *D. melanogaster,* we did not observe any major differences in LD decay within vs. between genes for either mutational class (Figure 3D, Supplementary Figure 5). In *C. grandiflora,* however, we observed significantly higher LD for within gene pairs of mutations compared to between genes pairs for both non-radical missense mutations and synonymous mutations (Figure 3C). Higher intra-gene LD was also evident if we calculated unsigned (*r^2^*) LD for *C. grandiflora* hinting at potential differences in recombination leading to faster LD decay between genes rather than LD created by epistasis (Supplementary Figure 6). Given that unsigned LD decay should mostly be driven by the rate of recombination, we hypothesize this difference in inter- vs intragenic LD is due to the strong enrichment of cross-overs in promoter regions of plant genomes (Choi et al. 2013; Hellsten et al. 2013). Such crossovers should rapidly erode LD between genes, while leaving within gene LD unaffected. Conversely, no such pattern is known to occur in flies where transcription start sites have actually been found to negatively correlate with cross-over occurrence (Comeron et al. 2012; Smukowski Heil et al. 2015).

**Figure 3.**
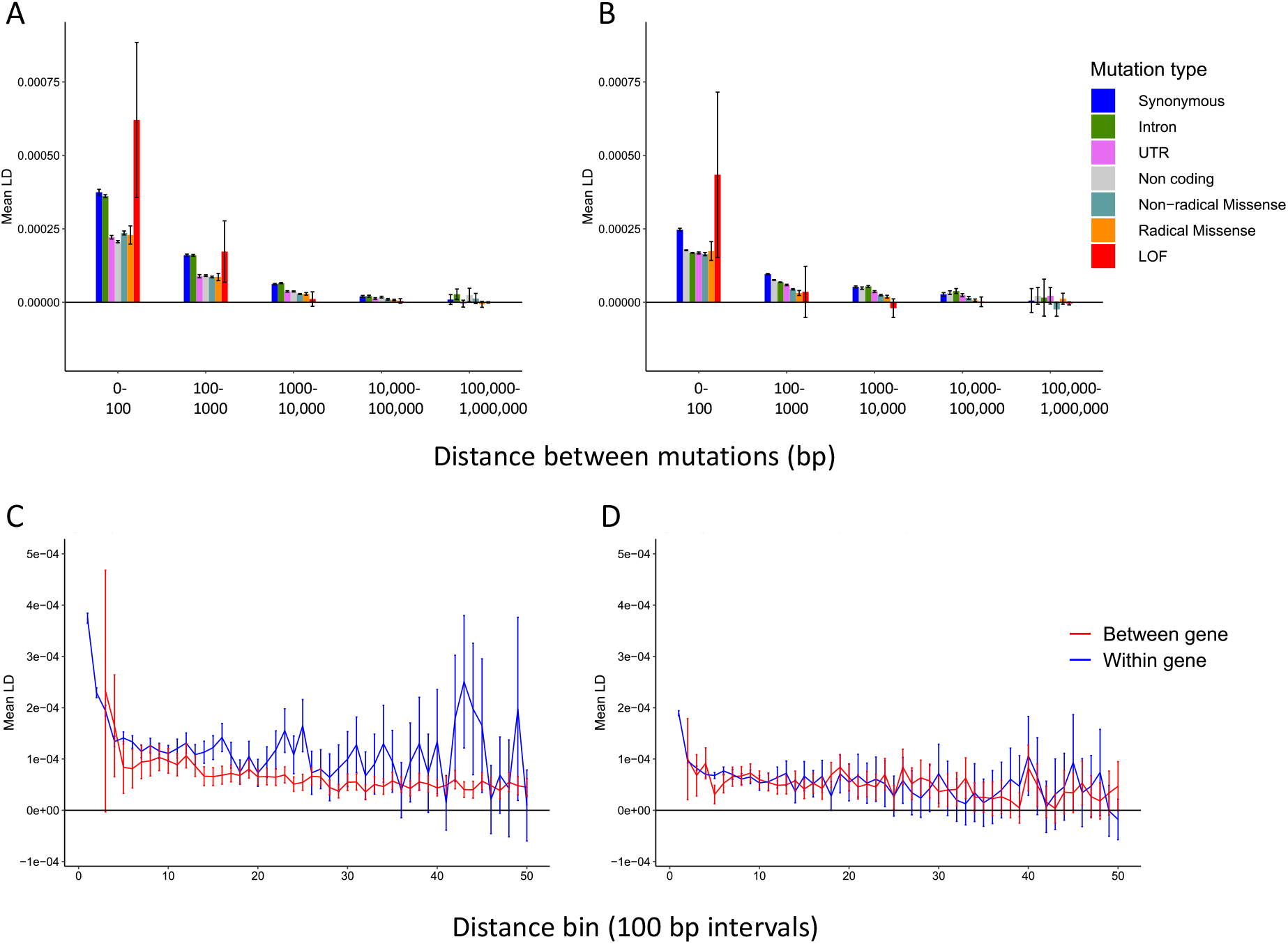
A,B. Distribution of mean signed LD for pairs of mutations across different distance bins for several mutation classes in A) *C. grandiflora* and B) *D. melanogaster*. Mutations within each bin are sorted by degree of expected deleteriousnes in ascending order. C,D Mean signed LD in 100bp bins for synonymous mutation pairs within and between genes for C) *C. grandiflora* and D) *D. melanogaster*.

Though our previous analyses found no obvious signature of pervasive intergenic synergistic epistasis when considering the entire genome, epistasis may be stronger between functionally related genes. To investigate this possibility, we examined LD among variants within interacting gene networks, using either radical missense or synonymous mutations. We obtained gene lists of metabolic and signalling networks from KEGG and only considered LD calculated among sites in different 100kb blocks to minimize any contribution of LD between nearby mutations and to remove measurements of LD between mutations within genes. We calculated the mean LD within each network, then averaged these mean LD values across networks (weighting by network size) to estimate “average network LD”. We permuted the assignment of genes to each KEGG network 1000x to create a null distribution for average network LD.

The mean LD of radical missense mutations within each network is shown in Figures 4A, B. Permutation tests indicated that the average network LD was significantly more negative than expected in *D. melanogaster* (average network LD = −5.71E-08, p = 0.008) but not in *C. grandiflora* (average network LD = −1.05E-08, p = 0.956). Figures 3A, B give the appearance that LD is related to network size but this is likely a statistical artifact that also occurs in permutations. To visualize this, we split networks into deciles with respect to network size and calculated average network LD for each decile. The permutation distributions had negative median values that approach zero for larger network sizes (Figures 3C, D). Overlaying the observed values on these permutation distributions helps visualize that the observed LD in *D. melanogaster* is more negative than expected across most network sizes (red points are empirical values and grey points are means from the distribution of permutation values). Repeating the permutation analysis with synonymous mutations, we found that average network LD was again significantly more negative than expected in *D. melanogaster* (average network LD = −1.03E-08, p = 0.006) but not *C. grandiflora* (average network LD = 9.89E-09, p = 0.902, see also Supplementary Figure 7 and Supplementary Table 2). Though the point estimate of LD is more strongly negative for radical missense than synonymous mutations in *D. melanogaster*, the qualitatively similar pattern complicates the interpretation (see Discussion).

**Figure 4.**
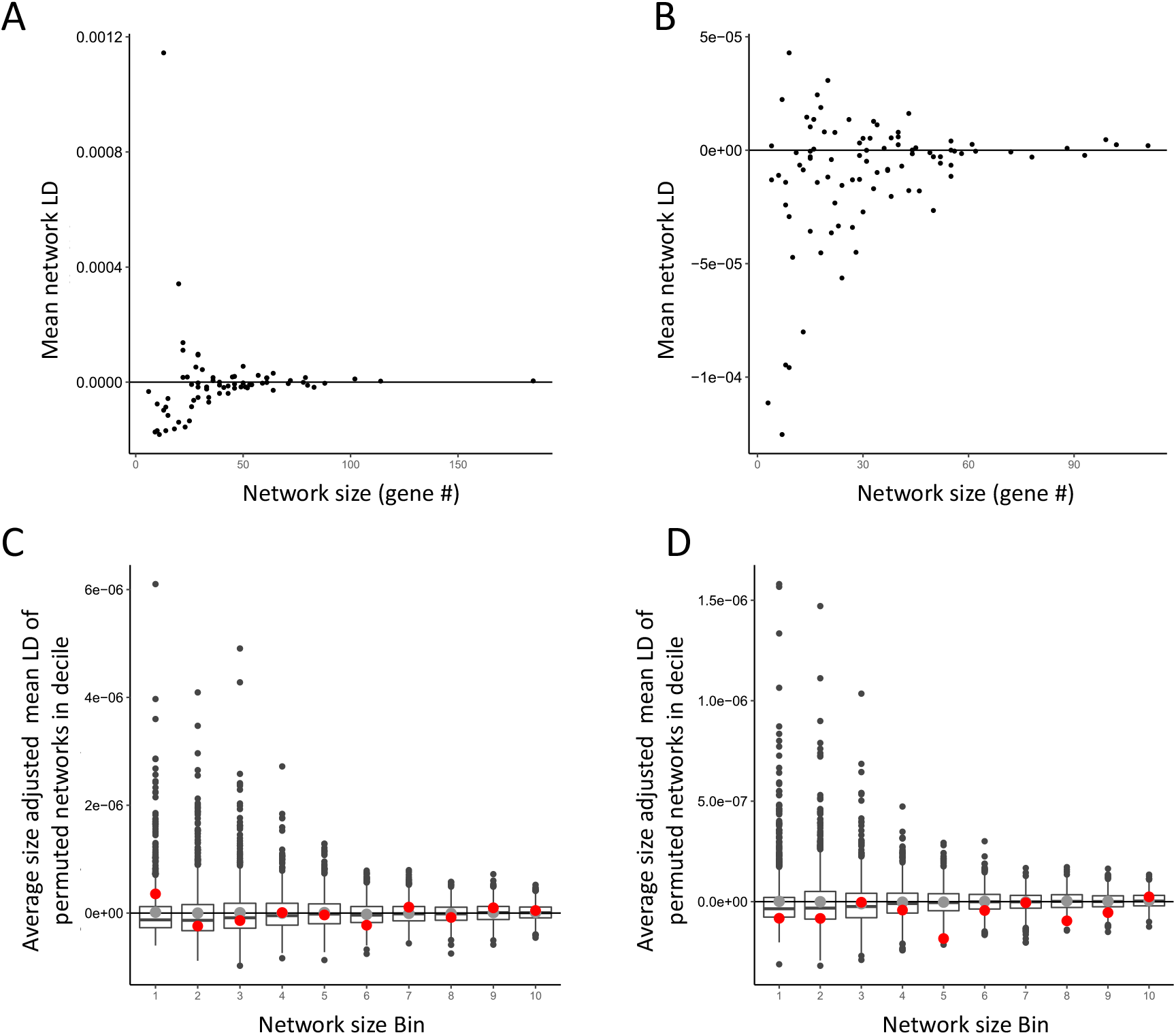
Mean LD among radical missense mutations affecting genes within interacting biological networks plotted against network size (as defined by numbers of genes within each network). LD was calculated among sites in different 100kb blocks to minimize the effects of short intra-genic interactions. Left data from *C. grandiflora*, right from *D. melanogaster.* C,D) Average network LD among radical missense mutations for deciles based on network size; LD values were weighted by network size. Box-plots show the null distribution for average network LD from permuted networks; black bar represents the median, the grey point represents the mean, and whiskers represent quartiles. In each permutation, networks were split into deciles based on bin size and the average LD of all networks in each decile was calculated. True average network LD of each decile is overlaid in red. Left data from *C. grandiflora*, right from *D. melanogaster*

## Discussion

In this study we analyzed patterns of signed LD in two species*, C. grandiflora* and *D. melanogaster*. When calculating mean LD among various classes of mutations, we found that less deleterious mutations tended to have more positive LD, with only LOF mutations exhibiting negative point estimates of mean LD under certain allelic-count cut-offs. Though the reduction in LD for deleterious classes such as LOF mutations relative to putatively neutral ones (e.g., synonymous mutations) could be interpreted as evidence of negative synergistic epistasis (Sohail et al. 2017) or Hill-Robertson interference (Garcia and Lohmueller 2020), other processes may provide more parsimonious alternatives. In particular, positive LD could be created by processes such as low level admixture in our datasets (Pfaff et al. 2001), and this effect may be weaker for more deleterious variants. For neutral sites, admixture can generate positive LD if LD is polarized either by rarity (as we have done) or by ancestral state if a minor allele threshold is imposed. Simulations across a range of demographic scenarios (both our own and those of Sohail et al.) have shown that positive LD builds up between mutations in a manner dependent on their selection coefficient; the more deleterious the mutations, the less positive LD builds up among them (Sohail et al. 2017), under a multiplicative model of negative selection. Presumably, the reason that positive LD occurs for low frequency neutral but less so for selected SNPs is as follows. Low frequency neutral SNPs within a given region will tend to be of two types: local variants of relatively recent origin but also migrant variants (of older origin), which will have come to the local population linked to migrant variants at other genomic sites (i.e., in positive LD). Deleterious variants are less likely to be of older (migrant) origin by virtue of the selection against them. Good (2020) showed that even without admixture, positive LD is expected between rare neutral mutations. This positive LD occurs because some variants that are rare in the present will have been more common in the past, providing an opportunity for a second variant to arise on the same haplotype. Positive LD is less likely to arise in this manner between deleterious variants because a deleterious variant is less likely to have been at higher frequency in the past. Though positive LD can arise in this fashion at both neutral and selected sites, admixture (including subtle forms of geographic structure) can potentially cause much stronger positive LD (Figure 2).

Part of our signal of genome-wide positive LD could be explained by the presence of multinucleotide mutations (Schrider et al. 2011; Ragsdale 2021). Multinucleotide mutations create strong positive LD among de-novo mutations, and such coupled mutations should persist much longer if both variants are neutral, potentially creating our observed pattern of an excess of positive LD for less deleterious mutations. However, previous work in humans has suggested that the majority of SNPs in multinucleotide mutations fall within 20bp of each other, which should create signed LD on a much smaller scale than what we have observed in our data (Schrider et al. 2011). Our genome-wide measures of LD are not driven exclusively by nearby sites; the LD measures are similarly positive even when we measure LD among sites in different 100kb blocks, thereby excluding the contribution of LD from the vast majority of neighbouring sites (Figure 1).

The fact that differences in LD between selected and neutral sites can arise in several simple models necessitates caution in interpretating differences in LD among mutation classes with varying deleteriousness. For example, previous studies have used LD among synonymous mutations as a control group for inferring synergistic epistasis (Sohail et al. 2017) or Hill-Robertson interference (Garcia and Lohmueller 2020). However, as outlined above, differences in LD for deleterious versus neutral mutations may be expected even under purely multiplicative selection, even without invoking selective interference.

Because of its importance in theoretical population genetics (Kimura and Maruyama 1966; Crow and Kimura 1970; Crow and Kimura 1979; Kondrashov 1982; Barton 1995), we were particularly interested in looking for evidence of synergistic epistasis in the form of negative LD at selected sites. Instead of comparing LD at selected and neutral sites, we used randomization tests to test whether negative LD among LOF mutations is significantly different from 0; it is not in either species. This approach is somewhat conservative, because processes like admixture may oppose the signal of negative LD created by synergistic epistasis. However, because the admixture effect should be minimal for the most deleterious classes of mutations, this may not pose a major limitation in searching for a signature of negative epistasis. The power of recombination to destroy associations built by selection is likely a much more severe limitation on synergistic epistasis—if it is common—creating a detectable signature on genome-wide LD.

An additional issue with estimating mean LD across all genes is that this averaging may hide meaningful variation. For example, epistasis between functional sites within a gene may be fundamentally different in strength and/or sign than intergenic epistasis. Moreover, physically close site pairs, which will often be intragenic, will be less affected by recombination’s power to destroy associations built by epistatic selection or Hill-Robertson interference. We visualised the distribution of LD among several classes of mutations in both datasets, split across bins of inter-mutation distance. We observed non-zero LD most readily for nearby mutations across all mutation classes, and in all cases it was significantly positive. Excluding the first distance bin in our analysis (1-100bp), the magnitude of positive LD present in each mutation class was predicted well by the expected deleteriousness of each type of mutation. This is pattern can be explained by the simple scenarios of positive LD build-up outlined above.

One notable deviation from the pattern of stronger positive LD for less deleterious mutation classes was that, in first distance bin, LOF mutations had the most positive point estimates of mean LD. We hypothesize that this pattern is due to within-gene antagonistic epistasis, which is to be expected if a single LOF mutation is indeed sufficient to knock out the function of a gene. This echoes similar findings from Puchta et al. 2016 who demonstrated that antagonistic epistasis within a yeast snoRNA was prevalent among large effect deleterious mutations occurring within conserved domains because such mutations effectively acted as LOF variants and thus did not impact fitness multiplicatively when combined with other deleterious mutations. Ragsdale (2021) showed that LD for missense mutations within human protein functional domains is significantly more positive than expected, also hinting at a potential signal of within gene antagonistic epistasis.

Aside from within-gene epistasis, epistatic interactions may be stronger or more frequent between mutations in functionally related genes. In particular, given that genes function as part of larger biological networks, negative epistasis may arise between deleterious mutations that affect the function of genes within the same networks (Chiu et al. 2012). To test this idea, we calculated mean LD among synonymous and among radical missense mutations present in genes within interacting biological networks defined by KEGG (Kanehisa et al. 2016). Permutation tests in *D. melanogaster* suggested that the observed intra-network LD among radical missense mutations was more negative than expected. Curiously, significantly negative network LD occurs for synonymous mutations too. This latter result is surprising for two reasons: (i) LD is (relatively) strongly positive for synonymous mutations at the genome-wide level (Figure 1), and (ii) negative epistasis should not affect (putatively neutral) synonymous sites. A possible explanation of these findings emerges from our suspicion that the overall genome-wide positive LD is due to processes of admixture and gene flow. The significantly negative network LD for both synonymous and radical missense mutations could be due to synergistic epistasis acting against introgressed alleles affecting the same network. Because introgressed haplotypes will include synonymous and missense mutations that are all in positive LD, selection on deleterious missense variants will lead to a drop in positive LD for multiple types of mutations.

Unlike *D. melanogaster*, network LD was not significantly negative in *C. grandiflora.* The lack of a significant result in *C. grandiflora* could be biologically meaningful or more mundane. For example, KEGG network delineation could be more biologically meaningful in *D. melanogaster* compared to *C. grandiflora* where network information has been obtained from a species in a different genus (*Arabidopsis thaliana*). Alternatively, the difference between species could simply be a statistical artifact (i.e., false positive in *D. melanogaster* or false negative in *C. grandiflora*). Similar analyses in other species will shed light on whether signed LD is related to network status.

Our examination of LD has revealed variation in the strength and, in some cases, the direction of signed LD. This variation is affected by several factors including proximity of sites, putative deleteriousness of mutations, and the functional relationship among genes. Some, but not all, of the patterns are consistent across two very different species. Some of these patterns can be generated by more than one process and, consequently, it will be challenging to conclusively prove which processes drive such patterns. Nonetheless, patterns of LD can serve as one line of evidence for (or against) particular hypotheses that are investigated using multiple approaches.

## Materials and Methods

### Population genomics datasets

We retrieved data from whole genome sequencing of 182 *C. grandiflora* individuals from (Josephs et al. 2015) and data for 197 haploid *D. melanogaster* embryos from the *Drosophila* population genomics project (DPGP3)(Lack et al. 2015). SNP calls previously generated by Josephs et al. for *C. grandiflora* were provided by Tyler Kent (personal communication). SNP calls for *D. melanogaster* were downloaded from the PopFly website (Hervas et al. 2017, http://popfly.uab.cat/). Both data sets are a result of thorough sampling from single populations with low population structure, making them ideal candidates for detecting signs of epistasis from patterns of LD. To ensure that recent migrants did not affect our LD analyses we used the R package SNPrelate (Li 2011) to visualize relatedness through PCA between *C. grandiflora* samples. This revealed six divergent genotypes that we eliminated from our downstream analysis leaving us with a total of 176 individuals. A previous study by Sohail et al. (2017) had already used the DPGP3 dataset to analyze patterns of LD so we used the 190 individuals they retained after their filtering in our own analyses. We further filtered both datasets by only considering bi-allelic sites where all individuals had genotype information. Following Sohail et al. (2017) we removed SNPs segregating within chemo-sensory and odorant binding genes in the *D. melanogaster* dataset based on gene lists obtained from FlyBase (Larkin et al. 2021), though their inclusion has little effect on the results. One final complication of the *D. melanogaster* dataset is the segregation of several large-scale inversions in this species. The initial establishment of an inversion creates some LD. However, gene exchange between chromosomes of different inversion karyotypes still occurs within inverted regions via double cross-over recombination events and gene conversion. Indeed, Houle and Márquez (2015) found that LD was only slightly stronger within versus outside LD regions. To the extent inversions cause a reduction in the effective recombination rate, inversions should amplify the ability to detect the existing signal of non-zero LD built by other forces (e.g., selection, migration). Nonetheless, we repeated the majority of our analyses excluding regions known to harbor inversions in the DPGP3 population. We obtained coordinates of such inversions from Corbett-Detig and Hartl (2012) and removed SNPs segregating in such regions for a subset of our analyses. However, analyses excluding inverted regions are necessarily based on much less data and consequently have reduced power.

### SNP annotation

We used SNPeff (Cingolani et al. 2012) and the genome annotations of the reference genomes (Slotte et al., 2013 for *Capsella rubella*; *D. melanogaster* release 5.57 from Thurmond et al., 2019) to functionally annotate SNPs in both datasets as either LOF, synonymous, missense (non-synonymous), intronic (but not splice affecting), UTR (if the SNP coordinate was either in the 5’ or 3’ UTR of a gene), or non-coding (for SNPs not present in coding regions). A small number of SNPs had annotations in multiple categories (e.g. both UTR and intronic), primarily due to multiple gene overlap, and were excluded from the analysis. We included stop-gain and splice-disrupting SNPs in our set of LOF mutations based on the method of Sohail et al. We also further classified missense SNPs as either radical or non-radical. Missense SNPs were considered radical if they changed both the volume and polarity of an amino acid based on previous work suggesting that change in either category lead result in particularly deleterious mutations in species such as *D. melanogaster* (Sainudiin et al. 2005; Weber and Whelan 2019, see also Supplementary Table 1 for the list of amino acid properties we used).

### Calculating LD

We calculated LD values in two ways. First, we used the same method as Sohail et al. by calculating a point estimate of average LD among all mutations. For a genome with *K* loci, let *X_i_* be a discrete, random variable representing the number of derived alleles present at locus *i*, which can take values 0, 1 for a haploid population or alternatively 0, 1, 2 for a diploid population. The variance in the total number of derived mutations carried by each individual in the population can be expressed as:

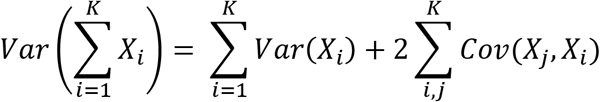

Because LD is, by definition, a covariance in the allelic state between two loci, we can use this equation to estimate the sum of all covariances across all loci by subtracting first term of the right-hand side from the term on the left-hand side (and then dividing by 2). The term on the left-hand side represents the genome-wide variance in mutation burden; the first term on the right-hand side is the sum of the variance in mutation burden at each locus. We can then estimate a mean value of LD per pair of loci by dividing by the number of possible two-way interactions in the dataset

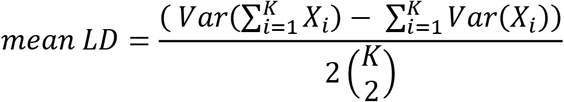

We also modified this approach to calculate LD on a block by block basis instead of SNP by SNP. This measure of average LD largely eliminates LD between physically close sites, which could initially arise via random mutation. We first split the genome into 100kb non-overlapping blocks. For a given genotype, we define *B_g_* as the number of derived variants in block *g*. This new variable can take values from 0 to 2*(number of segregating derived alleles in the given genomic block). To calculate total LD among all blocks, we infer the covariance in mutation burden between all blocks as follows

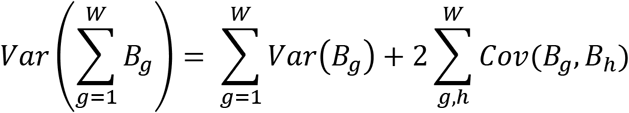

where *W* refers to the total number of 100kb blocks in the genome. Consider for example the simple case where we compare two blocks (*B*_*g*_, *B*_*h*_), each with two segregating sites, *B* can be represented as

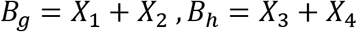

The number of covariance terms for these two genomic blocks is

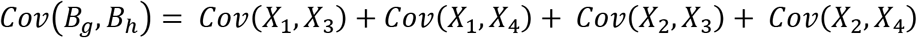

The within-block LD (e.g., *Cov*(*X*_1_, *X*_2_) and *Cov*(*X*_3_, *X*_4_)) from physically neighbouring sites contributes to the block-level variances (e.g., *Var*(*B*_*g*_) and *Var*(*B*_*h*_)) but not the between-block covariances. For an arbitrary number of blocks, *Cov*(*B*_*g*_, *B*_*h*_) can therefore be standardized per pair of interacting blocks as follows

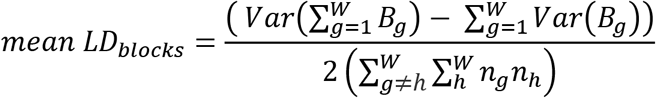

where *n*_*g*_ and *n*_*h*_ represent the number of sites with segregating derived variants in block *g* and *h* respectively.

We calculated mean LD using the above formula by transforming genotypes in our VCF files into tables of non-reference allele counts (0, 1, 2 for *C. grandiflora* and 0, 1 for *D. melanogaster*) and calculating the relevant statistics in R using the package matrixStats (Bengtsson 2017). We assumed that the non-reference alleles were the derived alleles in the two datasets. In principle, a reference genome assembled from a randomly sampled haplotype will contain some derived alleles that we will incorrectly assume are ancestral in our method. This issue however should be minimal since our analyses exclusively focus on rare mutations (<5% frequency) that are unlikely to be included in a reference assembly and will be filtered out as high frequency variants by our analysis even if they are included. This is especially true for most putatively deleterious mutations such as LOF mutations which are likely maintained at low frequency by mutation-selection balance.

We also calculated LD using PLINK (Purcell et al. 2007) for each category of mutation. We calculated LD using default PLINK parameters which involved subsampling LD observations as too many possible pairwise comparisons exist to reasonably compute the entire distribution of LD values for most classes of mutations. We estimated raw LD values by first estimating r between every single pair of mutations in our dataset in PLINK (using the *--r* option) and then back-calculating a raw value of LD by multiplying r by the square root of the product of allele frequencies at the two loci being compared. This approach allows us to observe the entire distribution of LD values rather than one summary statistic and back-calculating a raw value of LD from r allows us to compare values from our two methods directly. Finally, we binned distance between mutations pairs into seven categories: 100bp or less, 101-1000bp, 1001-10,000bp, 10,001-100,000bp, 100,001-1,000,000bp to visualize how signed LD decayed with distance for each class of mutations. Further, we compared signed LD decay within vs. between genes for synonymous and non-radical missense mutations. We did this by noting which gene our mutations of interest impacted according to SNPeff, and splitting our LD values into two categories, those where both contributing mutations occurred in the same gene, and those where both contributing mutations occurred in different genes. We then visualized LD as above, however, we only considered mutations 1-5000bp apart, and calculated mean LD in even 100bp bins, excluding any bins with less than 100 pairs of LD values.

### Gene network analysis

We used the R package Graphite (Sales et al. 2012) to obtain lists of genes from biological pathways described in the KEGG database (Kanehisa et al. 2016). Network information from KEGG was directly available for *D. melanogaster* but not *for C. grandiflora* where we instead used network information from *A. thaliana*. We used information on *C. grandiflora* - *A. thaliana* orthologs from (Josephs et al. 2015) to generate lists of interacting genes in *C. grandiflora*. Due to the low number of LOF mutations in each dataset we used low frequency (count of less than 5) radical missense mutations (definition described in SNP annotation section) as our set of candidate deleterious mutations. We calculated mean LD using 100kb blocks (as described above) for each network defined by KEGG for our two species, generating separating sets of networks for synonymous and radical mutations. Smaller networks (defined by the number of genes assigned by KEGG to each network) have more highly variable estimates of LD, presumably because of the smaller number of genes from which LD is estimated. Consequently, we calculated “size adjusted LD” values for all networks. We did this by correcting mean LD in 100kb blocks for each network as follows:

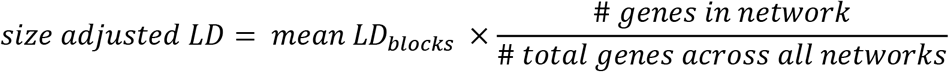

### Simulations

We used SLiM (V3.2.1) (Haller and Messer 2019) to run forward time simulations of population admixture to ask how signed LD can be affected by various demographic processes. We simulated three populations of 100,000 individuals each: one focal population that was sampled at the end of the simulation and two satellite populations with symmetrical migration to the focal population (10,000 individuals per generation). Each diploid individual in our simulation contained two 1Mb chromosomes with recombination and mutation rates both 1E-08 per bp per generation. Mutations were sampled from two categories: neutral (*s* = 0) with a probability 1%, or deleterious (*s* = −0.001) with a probability of 99%. Fitness was determined by the multiplicative effect of deleterious load in each individual genome, dominance was also assumed to be additive. We ran all simulations for 1.5 million generations altering the generation where continuous admixture was started in several treatment groups: no admixture, admixture starting at generation 200,000, 600,000, and 1,300,000. Each treatment group was made of 20 simulated replicates. After 1.5 million generations, we sampled 100 individuals from the focal population in each replicate. Next, we filtered out recent migrants in our focal population by performing a PCA on genotype of our samples and eliminating individuals with PC values greater than 1 SD away from the mean of PC1 or PC2. This mimics how we treated our real-world data where we eliminated outlier samples using PCA. Next, to replicate how we defined ancestral/derived alleles in the real-world data, we assigned all mutations with frequencies over 50% in our samples as the ancestral variant. Finally, we filtered out sites with a ‘derived’ allele count over 5 and calculated LD separately for neutral and deleterious mutations in each replicate. We also separately calculated LD for these simulations keeping the true ancestral state recorded by SLiM and polarizing LD by true ancestral/derived status both with and without a minor allele count cut-off, mimicking the way LD may be calculated in a real-world dataset where information on the true ancestral state may be available.

We ran a second set of simulations consisting of only one focal population where individuals were placed on a 2D landscape to simulate the effects of isolation by distance due to limited dispersal. We used the “Mate choice with a spatial kernel” recipe provided in the SLiM manual for this set of simulations. Briefly, 10,000 individuals were randomly placed on an (*x,y*) plane, with coordinate ranges [0,1] for both axes. To avoid clumping, individual fitness was calculated as a function of spatial competition with neighbouring individuals exerting the most costs to each other (see SLiM manual for more details URL: http://benhaller.com/slim/SLiM_Manual.pdf). Individuals chose mates a gaussian-distributed distance away, with mean 0, SD *σ*, and maximum value τ. We ran simulations with three sets of parameter values for *σ* and τ: (0.1,0.02), (0.3,0.06), (0.5,0.5). This range of values was selected to explore various levels of bias towards localized mating much like might occur in plant populations with limited pollen dispersal. Finally, offspring dispersed a gaussian distance away from their first parent. Each individual contained two 1Mb chromosomes containing only neutral mutations with a recombination and mutation rate of 1E-08 per bp per generation. The simulations were terminating after 100,000 generations and 100 individuals were sampled per simulation replicate. Each mate choice condition was replicated 10 times. After sampling, LD was calculated as described for the other simulations with the exception of PCA analysis as no migrant filtering was necessary due to the absence of cross-population migration.

## Data availability statement

Raw data for *C. grandiflora* are available at NCBI sequencing read archive (bio project ID: PRJNA275635). Data for *D. melanogaster* were downloaded via the PopFly website (Hervas et al. 2017, http://popfly.uab.cat/). Files containing annotated SNP calls (VCF format) used in this study will be made publicly available upon acceptance of this manuscript for publication. Scripts used in this study will also be made available at https://github.com/gsan211.

## Acknowledgements

The authors would like to thank Tyler Kent for helpful discussions and providing guidance on using *C. grandiflora* population genomic data. This work was supported by Natural Sciences and Engineering Research Council of Canada (NSERC) Discovery grants (S.I.W., and A.F.A.) and an NSERC Alexander Graham Bell Canada Graduate Scholarship (G.S.).

## Author Contributions

All authors designed the research. G.S. performed all analyses and wrote the draft manuscript. All authors revised the manuscript.

## Literature Cited

Agrawal AF, Whitlock MC. 2010. Environmental duress and epistasis: how does stress affect the strength of selection on new mutations? Trends Ecol. Evol. 25:450–458.

Auton A, McVean G. 2007. Recombination rate estimation in the presence of hotspots. Genome Res. 17:1219–1227.

Bank C, Hietpas RT, Jensen JD, Bolon DNA. 2015. A Systematic Survey of an Intragenic Epistatic Landscape. Mol. Biol. Evol. 32:229–238.

Barton NH. 1995. A general model for the evolution of recombination. Genet. Res. 65:123–145.

Chakraborty R, Weiss KM. 1988. Admixture as a tool for finding linked genes and detecting that difference from allelic association between loci. Proc. Natl. Acad. Sci. 85:9119–9123.

Chiu H-C, Marx CJ, Segrè D. 2012. Epistasis from functional dependence of fitness on underlying traits. Proc. R. Soc. B Biol. Sci. 279:4156–4164.

Choi K, Zhao X, Kelly KA, Venn O, Higgins JD, Yelina NE, Hardcastle TJ, Ziolkowski PA, Copenhaver GP, Franklin FCH, et al. 2013. *Arabidopsis* meiotic crossover hot spots overlap with H2A.Z nucleosomes at gene promoters. Nat. Genet. 45:1327–1336.

Cingolani P, Platts A, Wang LL, Coon M, Nguyen T, Wang L, Land SJ, Lu X, Ruden DM. 2012. A program for annotating and predicting the effects of single nucleotide polymorphisms, SnpEff. Fly 6:80–92.

Comeron JM, Ratnappan R, Bailin S. 2012. The Many Landscapes of Recombination in *Drosophila melanogaster*. PLOS Genet. 8:e1002905.

Corbett-Detig RB, Hartl DL. 2012. Population Genomics of Inversion Polymorphisms in *Drosophila melanogaster*. PLOS Genet. 8:e1003056.

Crow JF, Kimura M. 1970. An introduction to population genetics theory. Introd. Popul. Genet. Theory Ney York: Harper & Row

Crow JF, Kimura M. 1979. Efficiency of truncation selection. Proc. Natl. Acad. Sci. 76:396–399.

Elena SF, Lenski RE. 1997. Test of synergistic interactions among deleterious mutations in bacteria. Nature 390:395–398.

Good BH. 2020. Linkage disequilibrium between rare mutations. https://www.biorxiv.org/content/10.1101/2020.12.10.420042v1.full

Garcia JA, Lohmueller KE. 2020. Negative linkage disequilibrium between amino acid changing variants reveals interference among deleterious mutations in the human genome https://www.biorxiv.org/content/10.1101/2020.01.15.907097v1.full

Haller BC, Messer PW. 2019. SLiM 3: Forward Genetic Simulations Beyond the Wright– Fisher Model. Mol. Biol. Evol. 36:632–637.

Hellsten U, Wright KM, Jenkins J, Shu S, Yuan Y, Wessler SR, Schmutz J, Willis JH, Rokhsar DS. 2013. Fine-scale variation in meiotic recombination in *Mimulus* inferred from population shotgun sequencing. Proc. Natl. Acad. Sci. 110:19478–19482.

Hill WG, Robertson A. 1966. The effect of linkage on limits to artificial selection. Genet. Res. 8:269–294.

Hill WG, Robertson A. 1968. Linkage disequilibrium in finite populations. Theor. Appl. Genet. 38:226–231.

Houle D, Márquez EJ. 2015. Linkage Disequilibrium and Inversion-Typing of the *Drosophila melanogaster* Genome Reference Panel. G3 Genes Genomes Genet. 5:1695–1701.

Josephs EB, Lee YW, Stinchcombe JR, Wright SI. 2015. Association mapping reveals the role of purifying selection in the maintenance of genomic variation in gene expression. Proc. Natl. Acad. Sci. 112:15390–15395.

Kanehisa M, Sato Y, Kawashima M, Furumichi M, Tanabe M. 2016. KEGG as a reference resource for gene and protein annotation. Nucleic Acids Res. 44:D457–D462.

Kimura M, Maruyama T. 1966. The Mutational Load with Epistatic Gene Interactions in Fitness. Genetics 54:1337–1351.

Kondrashov AS. 1982. Selection against harmful mutations in large sexual and asexual populations. Genet. Res. 40:325–332.

Kondrashov AS. 1995. Dynamics of unconditionally deleterious mutations: Gaussian approximation and soft selection. Genet. Res. 65:113–121.

Lack JB, Cardeno CM, Crepeau MW, Taylor W, Corbett-Detig RB, Stevens KA, Langley CH, Pool JE. 2015. The Drosophila Genome Nexus: A Population Genomic Resource of 623 *Drosophila melanogaster* Genomes, Including 197 from a Single Ancestral Range Population. Genetics 199:1229–1241.

Lalić J, Elena SF. 2012. Magnitude and sign epistasis among deleterious mutations in a positive-sense plant RNA virus. Heredity 109:71–77.

Larkin A, Marygold SJ, Antonazzo G, Attrill H, dos Santos G, Garapati PV, Goodman JL, Gramates LS, Millburn G, Strelets VB, et al. 2021. FlyBase: updates to the Drosophila melanogaster knowledge base. Nucleic Acids Res. 49:D899–D907.

Li H. 2011. A statistical framework for SNP calling, mutation discovery, association mapping and population genetical parameter estimation from sequencing data. Bioinforma. Oxf. Engl. 27:2987–2993.

McEvoy BP, Powell JE, Goddard ME, Visscher PM. 2011. Human population dispersal “Out of Africa” estimated from linkage disequilibrium and allele frequencies of SNPs. Genome Res. 21:821–829.

McVean G. 2007. The Structure of Linkage Disequilibrium Around a Selective Sweep. Genetics 175:1395–1406.

Pfaff CL, Parra EJ, Bonilla C, Hiester K, McKeigue PM, Kamboh MI, Hutchinson RG, Ferrell RE, Boerwinkle E, Shriver MD. 2001. Population structure in admixed populations: effect of admixture dynamics on the pattern of linkage disequilibrium. Am. J. Hum. Genet. 68:198–207.

Puchta O, Cseke B, Czaja H, Tollervey D, Sanguinetti G, Kudla G. 2016. Network of epistatic interactions within a yeast snoRNA. Science 352:840–844.

Purcell S, Neale B, Todd-Brown K, Thomas L, Ferreira MAR, Bender D, Maller J, Sklar P, de Bakker PIW, Daly MJ, et al. 2007. PLINK: A Tool Set for Whole-Genome Association and Population-Based Linkage Analyses. Am. J. Hum. Genet. 81:559–575.

Ragsdale AP. 2021. Can we distinguish modes of selective interactions using linkage disequilibrium? bioRxiv:2021.03.25.437004.

Sainudiin R, Wong WSW, Yogeeswaran K, Nasrallah JB, Yang Z, Nielsen R. 2005. Detecting Site-Specific Physicochemical Selective Pressures: Applications to the Class I HLA of the Human Major Histocompatibility Complex and the SRK of the Plant Sporophytic Self-Incompatibility System. J. Mol. Evol. 60:315–326.

Sales G, Calura E, Cavalieri D, Romualdi C. 2012. graphite - a Bioconductor package to convert pathway topology to gene network. BMC Bioinformatics 13:20.

Schrider DR, Hourmozdi JN, Hahn MW. 2011. Pervasive Multinucleotide Mutational Events in Eukaryotes. Curr. Biol. 21:1051–1054.

Slotte T, Hazzouri KM, Ågren JA, Koenig D, Maumus F, Guo Y-L, Steige K, Platts AE, Escobar JS, Newman LK, et al. 2013. The *Capsella rubella* genome and the genomic consequences of rapid mating system evolution. Nat. Genet. 45:831–835.

Smukowski Heil CS, Ellison C, Dubin M, Noor MAF. 2015. Recombining without Hotspots: A Comprehensive Evolutionary Portrait of Recombination in Two Closely Related Species of Drosophila. Genome Biol. Evol. 7:2829–2842.

Sohail M, Vakhrusheva OA, Sul JH, Pulit SL, Francioli LC, Consortium G of the N, Initiative ADN, Berg LH van den, Veldink JH, Bakker PIW de, et al. 2017. Negative selection in humans and fruit flies involves synergistic epistasis. Science 356:539–542.

Stephens JC, Briscoe D, O’Brien SJ. 1994. Mapping by admixture linkage disequilibrium in human populations: limits and guidelines. Am. J. Hum. Genet. 55:809–824.

Thurmond J, Goodman JL, Strelets VB, Attrill H, Gramates LS, Marygold SJ, Matthews BB, Millburn G, Antonazzo G, Trovisco V, et al. 2019. FlyBase 2.0: the next generation. Nucleic Acids Res. 47:D759–D765.

Weber CC, Whelan S. 2019. Physicochemical Amino Acid Properties Better Describe Substitution Rates in Large Populations. Mol. Biol. Evol. 36:679–690.

