## Supplementary Figures and Tables for "Patterns and Causes of Signed Linkage Disequilibria in Flies and Plants"

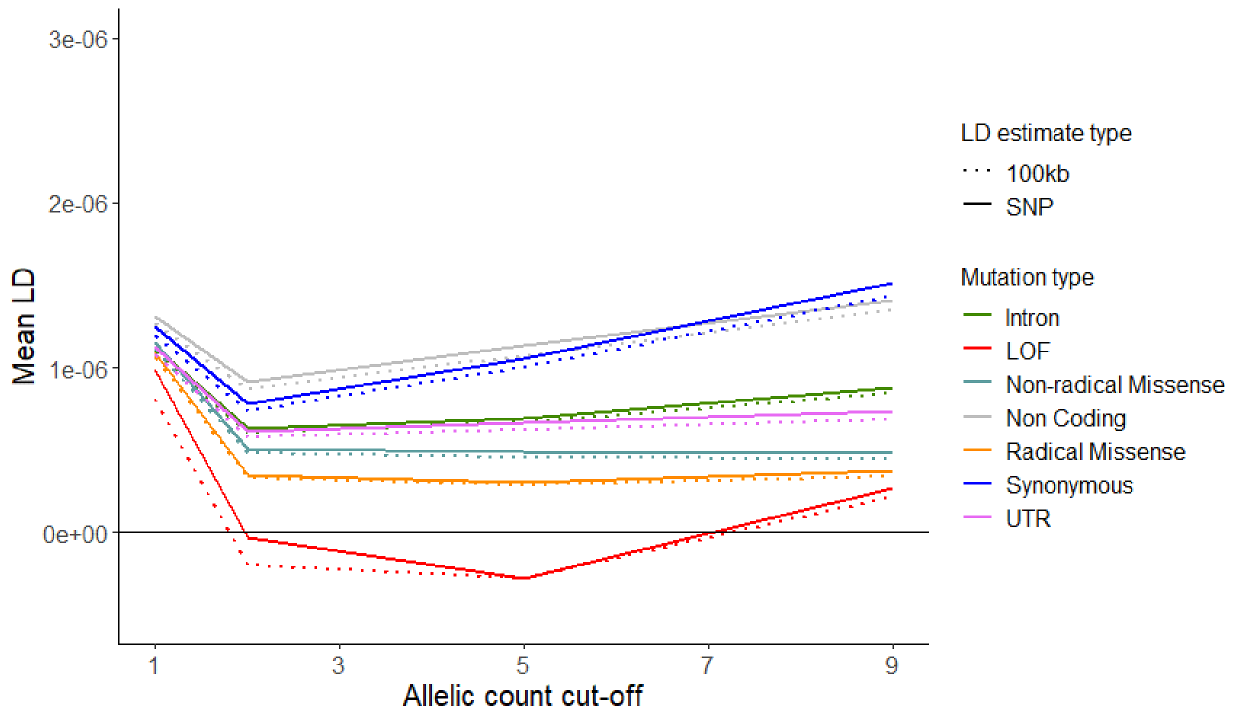


Supplementary Figure 1. Mean pair-wise LD among several classes of mutations across different allele count cut-offs. Data from *D. melanogaster* excluding regions harboring inversions. Solid lines indicate mean LD among all SNPs, dashed lines indicate LD calculated among sites in different 100kb, non-overlapping genomic blocks.

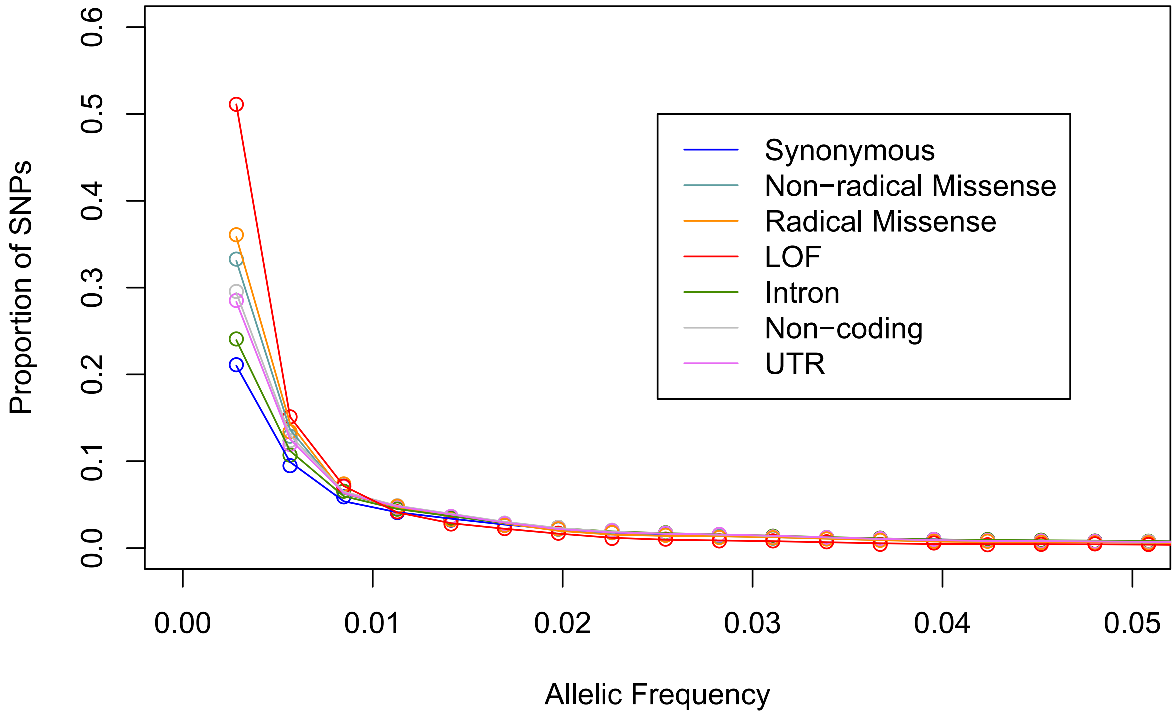


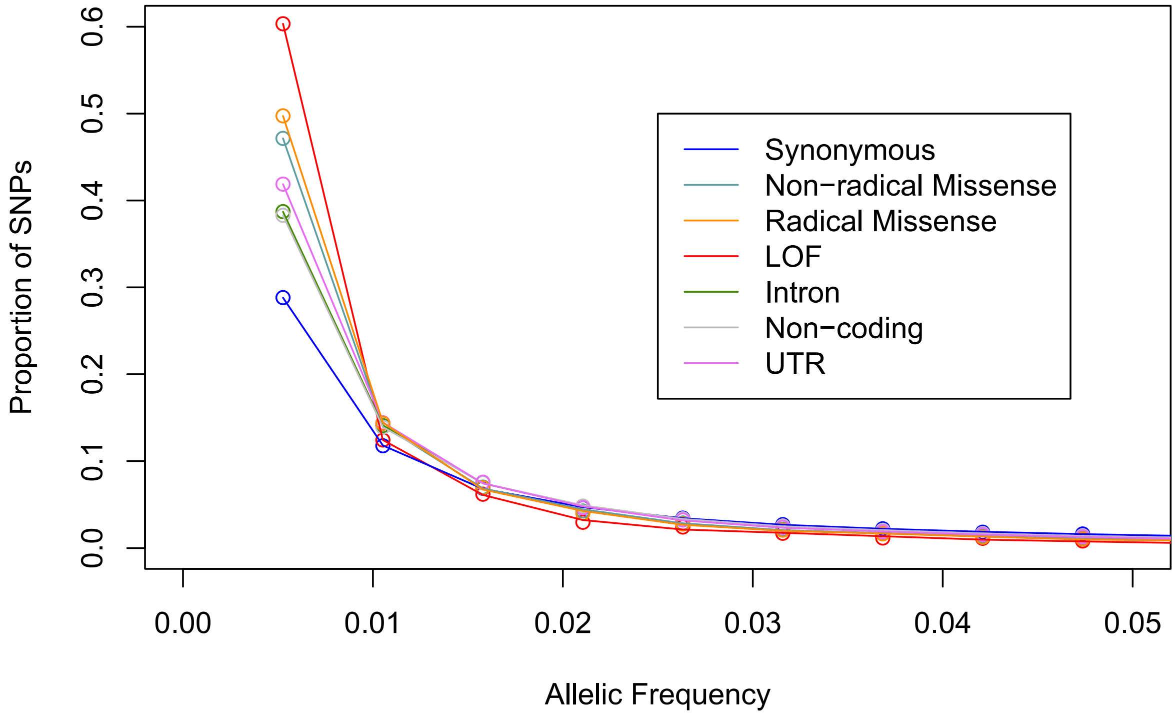


Supplementary Figure 2. Site frequency spectra for several classes of mutations. Top *C. grandiflroa*, bottom *D. melanogaster*.

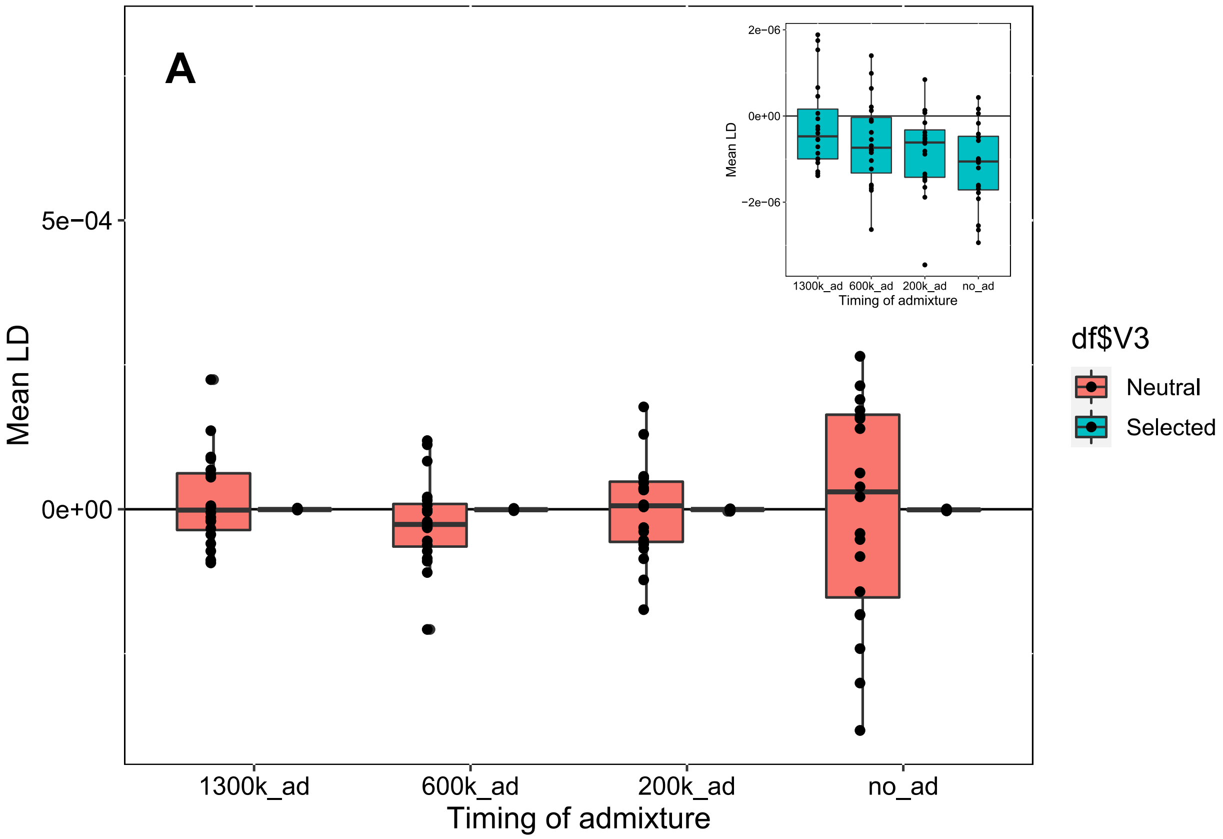
A
Mean LD
5e−04
0e+00
1300k_ad
600k_ad
200k_ad
no_ad
Timing of admixture
2e−06
0e+00
−2e−06
df$V3
Neutral
Selected

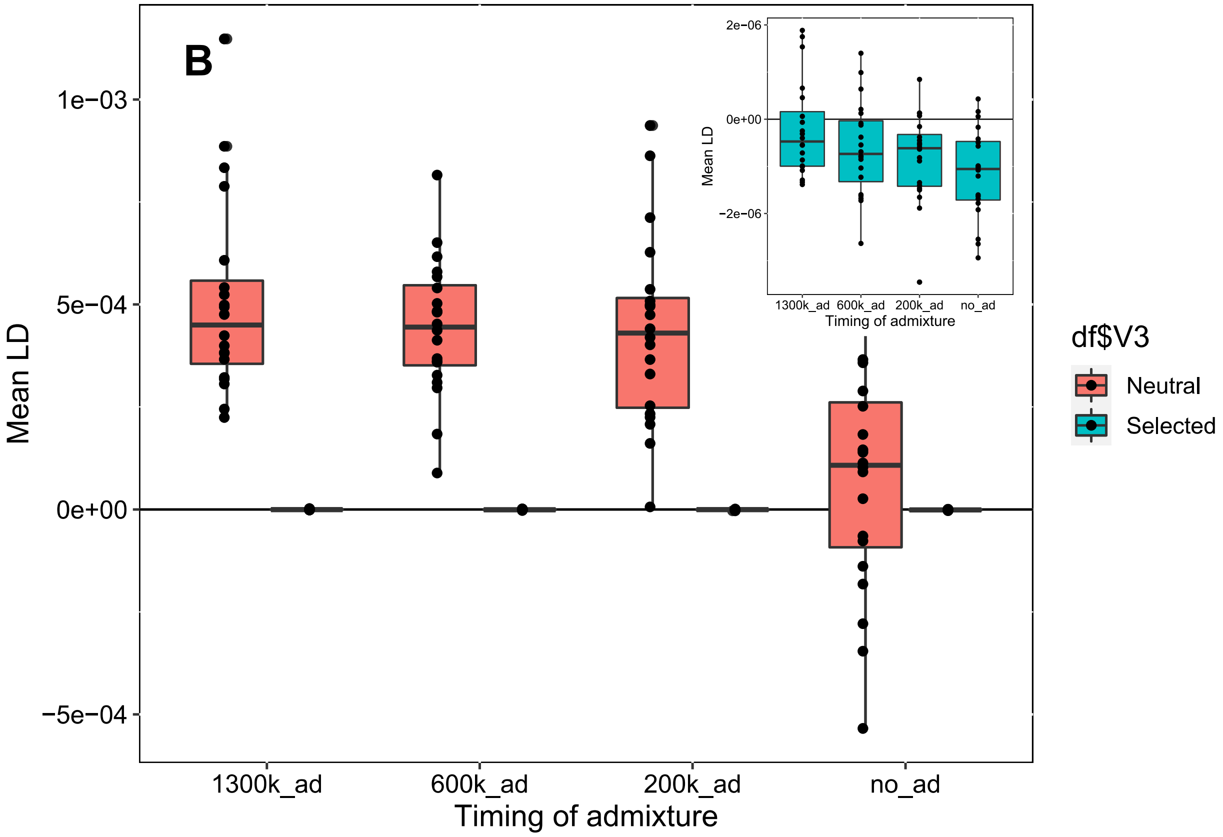
B
Mean LD
1e−03
5e−04
0e+00
−5e−04
1300k_ad
600k_ad
200k_ad
no_ad
Timing of admixture
2e−06
0e+00
−2e−06
df$V3
Neutral
Selected

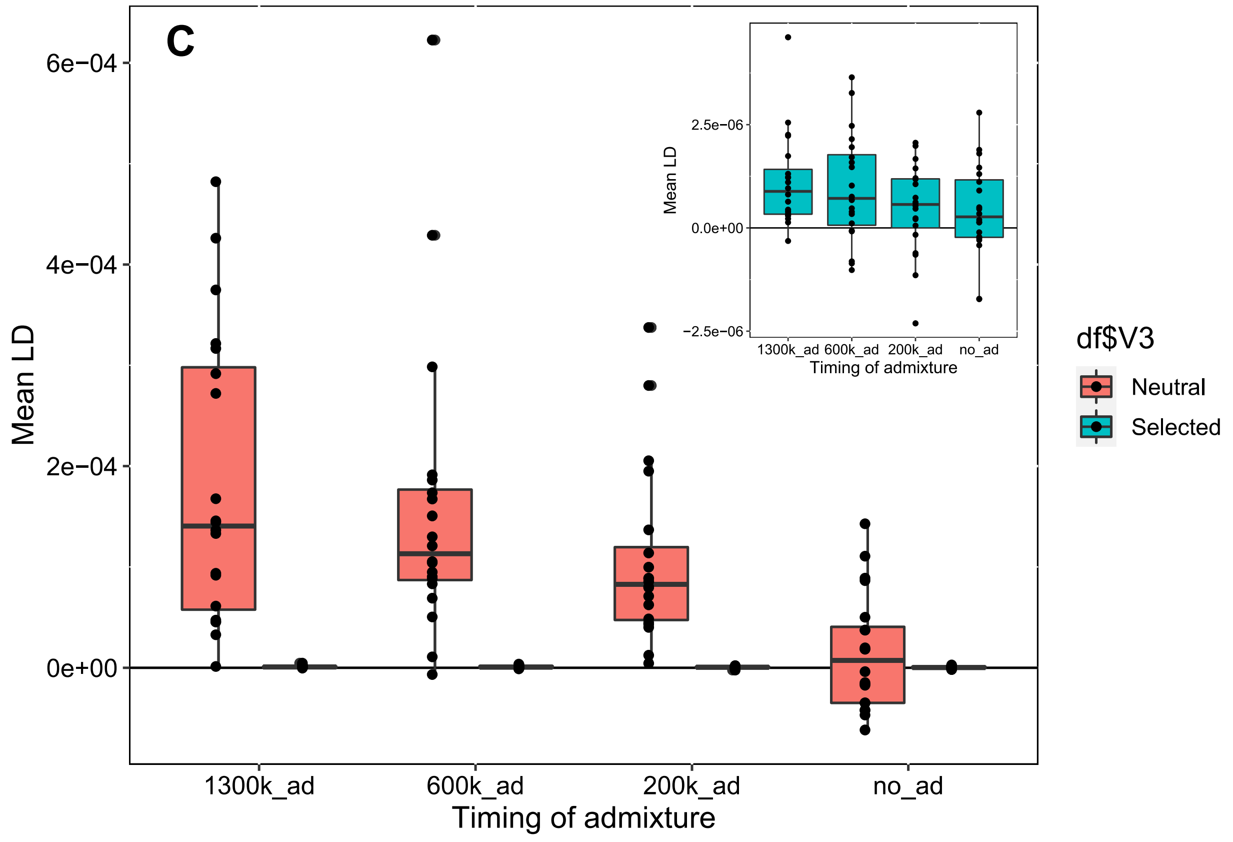
C
6e−04
4e−04
2e−04
0e+00
Mean LD
1300k_ad
600k_ad
200k_ad
no_ad
Timing of admixture
2.5e−06
0.0e+00
−2.5e−06
Mean LD
Timing of admixture
df$V3
Neutral
Selected

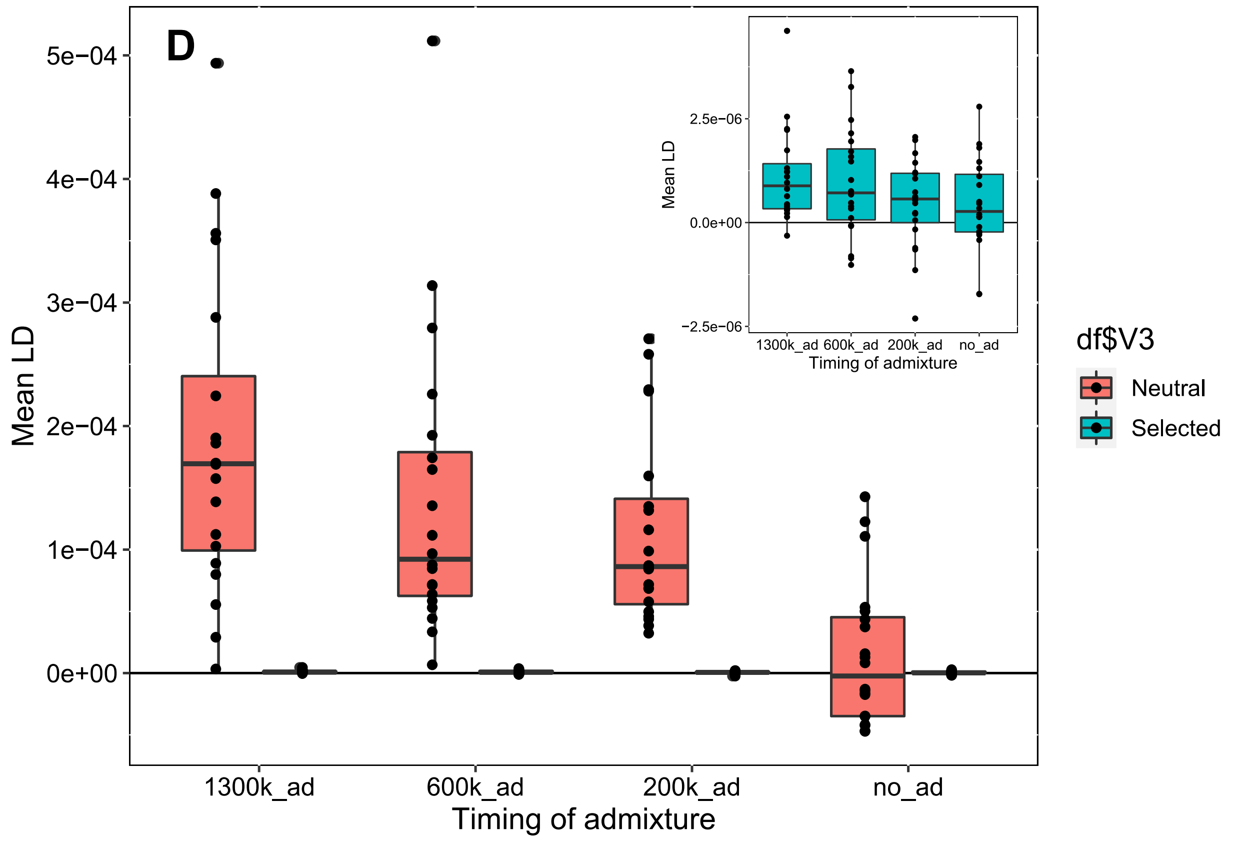
D
5e−04
4e−04
3e−04
2e−04
1e−04
0e+00
Mean LD
1300k_ad
600k_ad
200k_ad
no_ad
Timing of admixture
2.5e−06
0.0e+00
−2.5e−06
Mean LD
Timing of admixture
df$V3
Neutral
Selected

Supplementary Figure 3. Mean signed LD among simulated neutral and deleterious mutations under different scenarios of admixture. The $x$-axis represents the generation in which admixture between isolated populations started. All simulations were run of a total of 1.5 million generations. Inset highlights results for selected (deleterious) mutations. A,B) LD polarized by ancestral state, A) all mutations considered, B) only rare mutations considered. C,D) LD polarized by allelric rarity, C) all mutations considered, D) only rare mutations considered (copy of Figure 2A in main text).

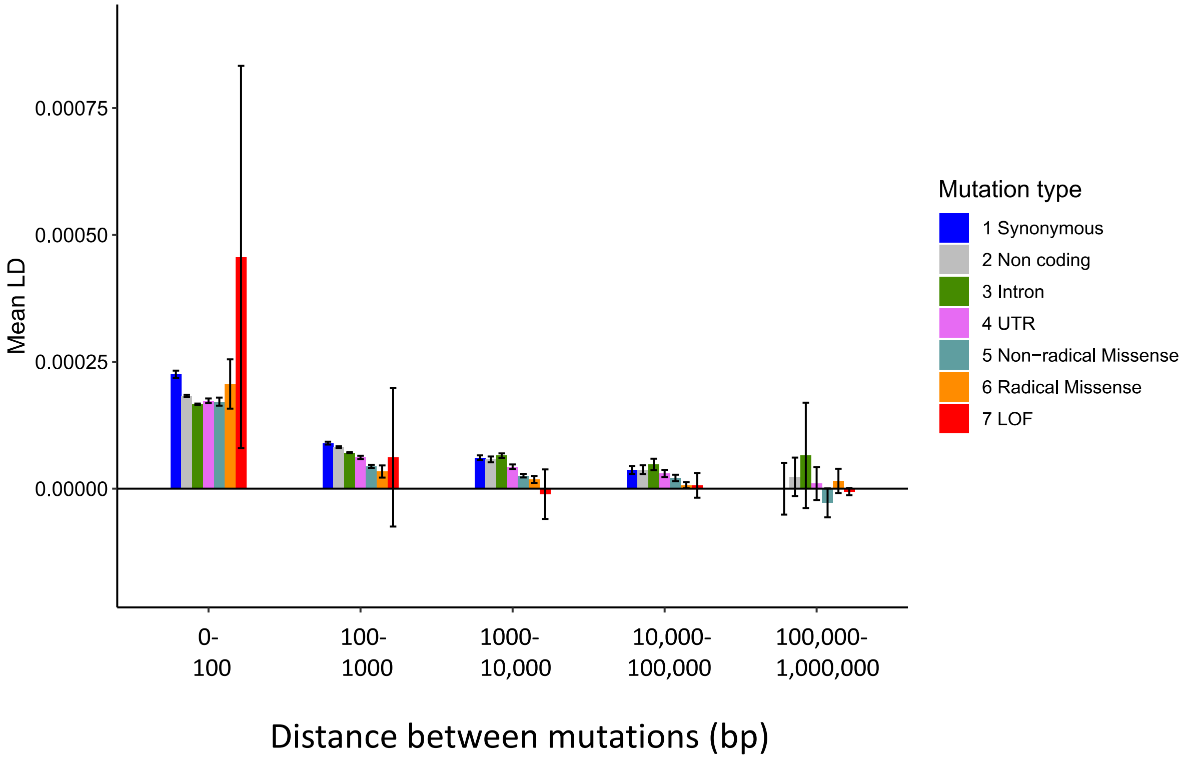


Supplementary Figure 4. Distribution of mean signed LD for pairs of mutations across different distance bins for several mutation classes. Data from *D. melanogaster* excluding regions harboring inversions.

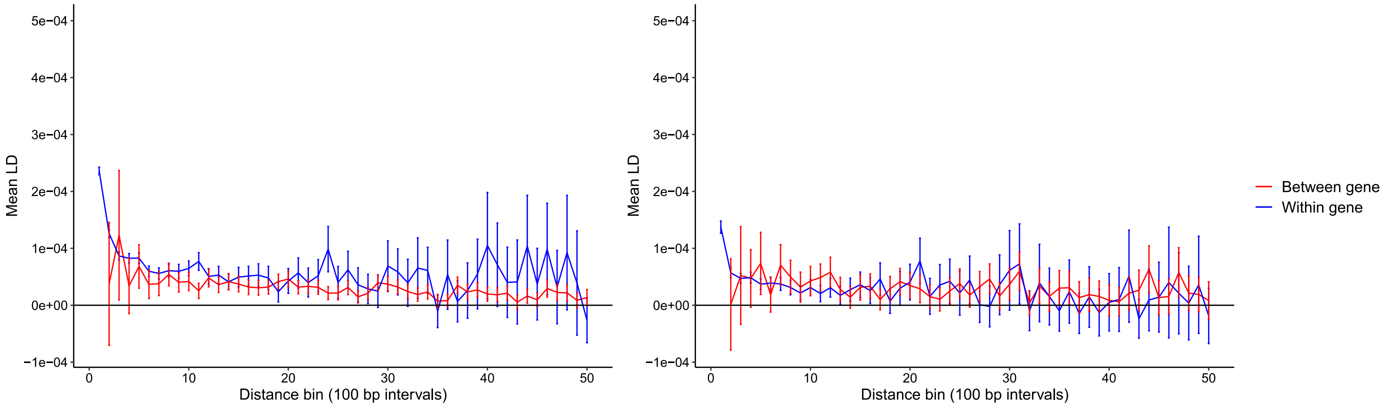


Supplementary Figure 5. LD decay within vs. between genes for non-radical missense mutations in *C. grandiflora* left and *D. melanogaster* right.

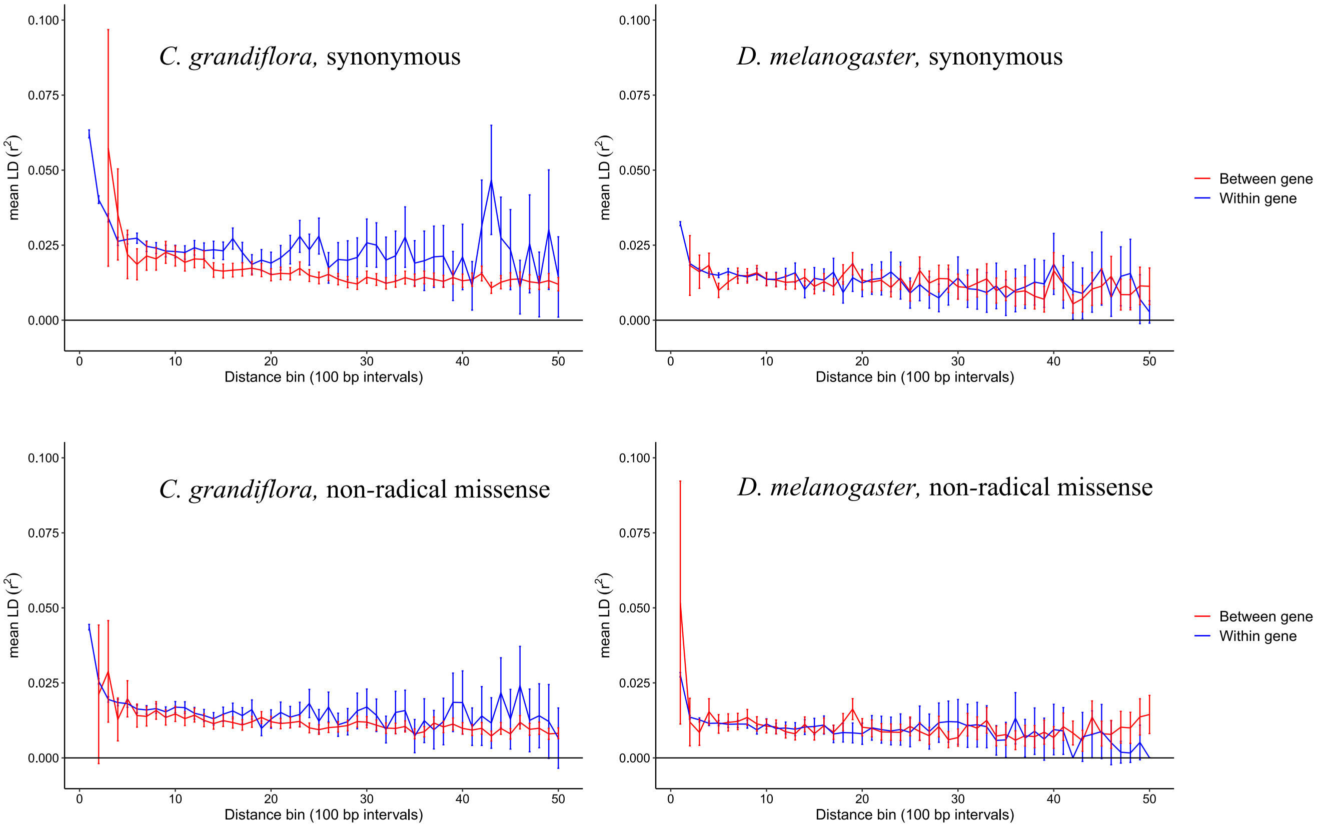


Supplementary Figure 6. Unsigned LD decay within vs. between genes in *C. grandiflora* left and *D. melanogaster* right. Top synonymous mutations, bottom non-radical missense mutations.

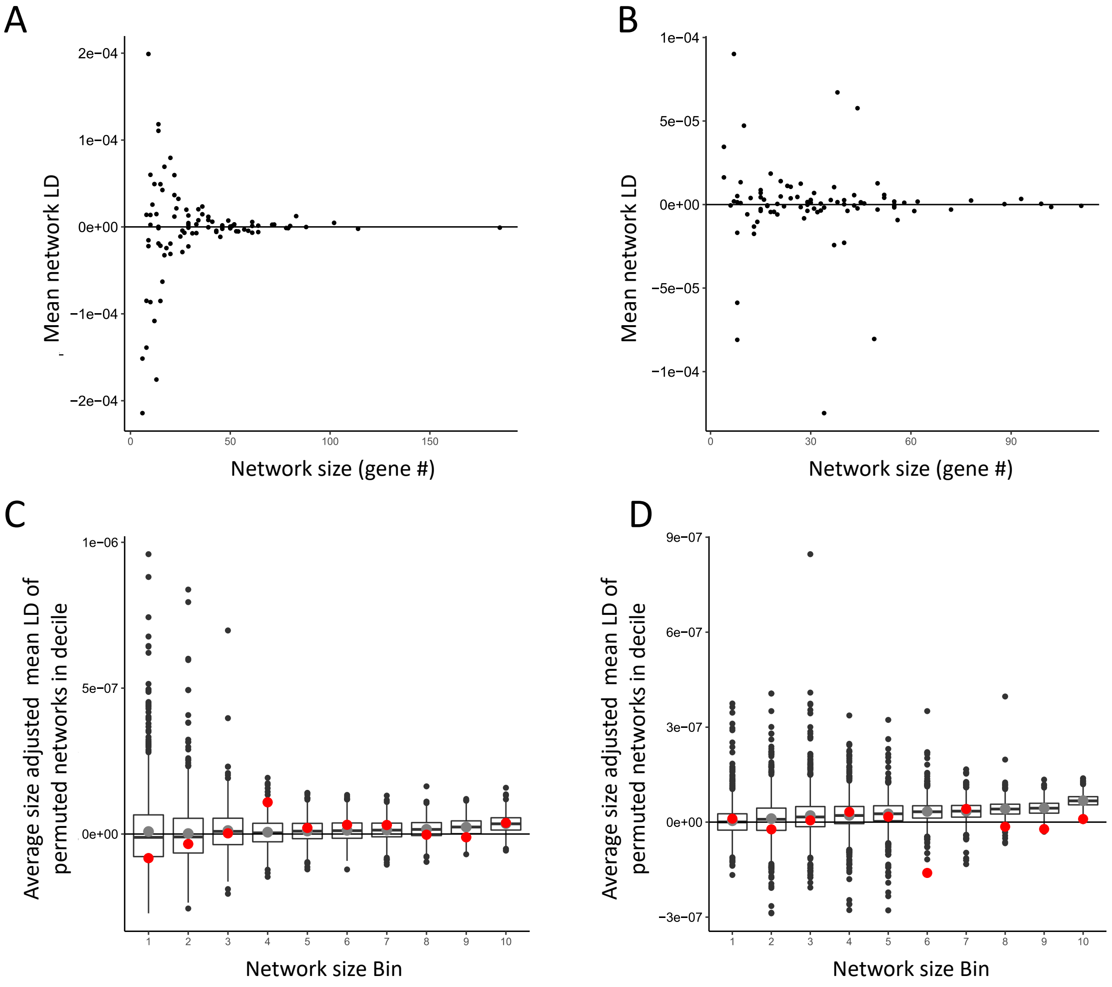


Supplementary Figure 7. Mean LD among synonymous mutations affecting genes within interacting biological networks plotted against network size (as defined by numbers of genes within each network). LD was calculated among sites in different 100kb blocks to minimize the effects of short intra-genic interactions. Left data from *C. grandiflora*, right from *D. melanogaster*. C,D) Average network LD among radical missense mutations for deciles based on network size; LD values were weighted by network size. Box-plots show the null distribution for average network LD from permuted networks; black bar represents the median, the grey point represents the mean, and whiskers represent quartiles. In each permutation, networks were split into deciles based on bin size and the average LD of all networks in each decile was calculated. True average network LD of each decile is overlaid in red. Left data from *C. grandiflora*, right *from D. melanogaster.*

Supplementary Table 1. List of amino acid properties used to define radical and non-radical missense changes.

| Amino Acid | Polarity | Size |
|---|---|---|
| Tyr | Polar | Large |
| Trp | Polar | Large |
| Thr | Polar | Small |
| Ser | Polar | Small |
| Arg | Polar | Large |
| Gln | Polar | Large |
| Asn | Polar | Small |
| Lys | Polar | Large |
| His | Polar | Large |
| Glu | Polar | Large |
| Asp | Polar | Small |
| Cys | Polar | Small |
| Val | Non-polar | Small |
| Pro | Non-polar | Small |
| Met | Non-polar | Large |
| Leu | Non-polar | Large |
| Ile | Non-polar | Large |
| Gly | Non-polar | Small |
| Phe | Non-polar | Large |
| Ala | Non-polar | Small |
